# Energy Metabolizes in Male Bone Marrow Mesenchymal Stem Cells Aging Process

**DOI:** 10.64898/2026.06.17.732798

**Authors:** Yue Chen, Huiyu Wang, Xin Lu, Jianxiang Zhao, Lijun Yang, Yadong Wang

## Abstract

Senescence human bone marrow mesenchymal stem cells (BMSCs), vulnerable to age-related defects, is poor in tissue regeneration. Cells in bone marrow accumulated senescent contributing to the development of metabolic energy regulation hold prospects for therapeutic advances. This study aimed to evaluate energy metabolic changes in male bone marrow mesenchymal stem cells senescence process. Our research established cell specific surface marker and enzymes expression level changes, as well as ECAR and OCR resonance. Notably, CD14, HLA-DRB1 and CD90 upregulated, glycolysis-related genes are increased, tricarboxylic acid cycle-related genes are decreased. We firstly identified links between time-dependent cell aging process and energy metabolism in BMSCs.

## Introduction

Human bone marrow mesenchymal stem cells (BMSCs) are adult stem cells in bone marrow, has the ability to be differentiated into osteoblast, chondrocyte, adipocyte under conditional medium (Chen et al. 2022, Yu et al. 2022).

Mesenchymal stem cells are a kind of plastic adherent cells, positive expression of CD73 (Sperber et al. 2023), CD90 (Moraes et al. 2016), CD105 (Greiner et al. 2022), and lack the expression of CD14, CD34, CD45, CD79A, and HLA-DR: CD14: -, CD14 molecule, be in placenta, liver, appendix, derived innate immune response to bacterial lipopolysaccharide, and to viruses.; CD34: -, CD34 molecule, be in placenta, fat, lung, mediated cell adhesion in bone marrow.; CD45: -, protein tyrosine phosphatase receptor type C, be in lymph node, appendix, spleen, may play partially roles in cell growth, differentiation, mitosis, and oncogenic transformation.; CD79A: -, CD79a molecule, be in lymph node, spleen, appendix, Ig-alpha protein of the B-cell antigen component, mediated B-cell function.; HLA-DR: -, major histocompatibility complex, class II, DR, major in cellular interaction during antigen presentation to CD4-positive T cells.; CD73: +, 5’-nucleotidase ecto, be in endometrium, ovary, colon, catalyze the conversion of extracellular nucleotides to membrane-permeable nucleosides.; CD90: +, Thy-1 cell surface antigen, be in brain, kidney, urinary bladder, involve in cell adhesion and communication, especially in immune system, as a marker be shared with hematopoietic stem cells.; CD105: +, endoglin, be in spleen, lung, ovary, mutations in the gene cause hereditary hemorrhagic telangiectasia.

In all, CD14 participates in inflammatory responses through the Toll-like receptor 4 (TLR4) signaling pathway, and chronic inflammation is closely associated with metabolic diseases such as metabolic syndrome, obesity, and diabetes. CD14 may influence insulin sensitivity and energy expenditure in adipose tissue by regulating the production of inflammatory cytokines. In obesity and diabetes, the imbalance in M1/M2 macrophage polarization may lead to abnormal energy metabolism. CD14 plays a crucial role in macrophage activation and polarization, potentially regulating energy metabolism by influencing this process.

The role of HLA-DRB1 in antigen presentation may affect the metabolic state of CD4+ T cells. For example, the metabolic reprogramming of Treg cells may be influenced by HLA-DRB1, thereby regulating systemic energy metabolism. Certain autoimmune diseases (e.g., rheumatoid arthritis) are associated with metabolic abnormalities, and specific polymorphisms of HLA-DRB1 may participate in the pathogenesis of these diseases by affecting immune-metabolic pathways.

The expression of CD90 in mesenchymal stem cells (MSCs) may influence their role in tissue repair and metabolic regulation. The metabolic activity of MSCs may be regulated through CD90, affecting the metabolic environment of local tissues.

Here we use in vitro cultured BMSCs, and zymogenic method combine with metabolites, associated with mitochondrial function, illustrated the energic phenotype of cell in vitro senescence. P7 and P13 cells were identified targeted metabolomics, gene expression, oxidative phosphorylation and glycolytic capacity prolonged with passage.

## Materials and methods

### Ethics statement

Ethical approval of the use of BMSCs cells was obtained from the ethical committee at Harbin Institute of Technology. We conducted this study in strict accordance with the regulation of 1964 Declaration of Helsinki and its later amendments of comparable ethical standards.

### Reagents and chemicals

Methanol and acetonitrile of HPLC grade were obtained from Fisher Scientific (Waltham, MA). formic acid and ammonium hydroxide were provided by Fluka (St. Louis, MO). Isopropanol and metayl tert-butyl ether (MTBE) were purchased from MREDA (Beijing, China). Deionized water was generated by a Milli-Q water purification system (Millipore, Billerica, MA).

### Cell culture

Two male BMSCs were obtained from Cyagen | OriCell™, cat# HUXMA-01001 (Guangzhou, China). The cells were cultured in complete medium for human bone marrow mesenchymal stem cells, OriCell™, cat#HUXMA-90011, following the user manual. Briefly, cell culture medium was refreshed every 3 days, and the cell density reached 80% for passage.

### Cellular Bioenergetic Function Analysis

Glycolytic or mitochondrial oxidative phosphorylation rates of p7 and p9 cells were measured by Seahorse XF24 analyzer. p7 and p9 cells were seeded at 30000 cells per well in 24-well overnight. In the OCR experiment, following overnight, p7 and p9 cells were washed and equilibrated at 37°C for 1h with the assay medium containing 10mM Glucose, 1mM Pyruvate and 2mM L-glutamine (Sigma-Aldrich, German). With the treatment of oligomycin, trifluorocarbonylcyanide phenylhydrazone (FCCP), rotenone and antimycin A (Seahorse Biosciences, UK) at 37°C for 20mins, the levels of mitochondrial oxidative phosphorylation were detected by seahorse XF24 analyzer. Oligomycin is an inhibitor of ATP synthase, which indirectly reflects the ATP production of cells. FCCP is an uncoupling agent, which evaluate the maximal respiratory capacity. Rotenone and antimycin A, both respiratory chain inhibitors, completely prevent mitochondrial oxygen consumption. In the ECAR experiment, p7 and p9 cells were washed and equilibrated at 37°C for 1h with the assay medium containing 2mM L-glutamine. Then treated with glucose, oligomycin and 2-DG (Seahorse Biosciences, UK), the glycolytic capacity of p7 and p9 cells were measured by seahorse analyzer. The addition of glucose represented the glycolytic ability of the cell at this time. After adding oligomycin, oxidative phosphorylation was inhibited. At this time, the cell completely depended on glycolysis for oxygen. The increase of acid production represented the glycolysis capacity. The total value represented the maximum glycolysis capacity. Finally, the glycolysis inhibitor 2-DG was added. The value after 2-DG showed that it was completely caused by acid production by a mechanism other than glycolysis.

### UPLC−MS Analysis

Metabolomic analysis was performed upon an ultra-high-performance liquid chromatography system (Shimadzu, Kyoto, Japan) coupled to a TripleTOF 5600+ series quadrupole time-of fight mass spectrometer (AB Sciex, Redwood City, CA). The chromatographic separation was performed by using an UPLC HSS T3 (100 × 2.1 mm, 1.8 μm) column (Waters, Milford, MA). The column temperature was set to 40°C during the analyses, and the injection volume was 5μL. A linear gradient elution with a total run time of 25 min was performed at a flow rate of 0.3mL/min with mobile phase A (isopropanol/methanol,9/1 v/v with 10mM ammonium formate) and B (acetonitrile water solution with 10mM ammonium formate) as follows: 10% A for 1.0 min;10-80% A from 1.0 to 15.0 min; 80-95% A from 15.0 to 20.0 min; holding 95% A for 4 min; a 1 min gradient returned to 10% A from 24.0 to 25.0 min; holding 2min to totally column recovery. Full scan and MS/MS scan were performed both in positive and negative electrospray ionization modes (ESI+, ESI-). The conditions were set as follows: spray voltages 5.5 kV in positive mode and 5.0 kV in negative mode; mass scan from 300 to 1500 m/z at a rate of 1.5 spectra/s; collision energy from 20 to 70 eV.

### Data Processing

Data pre-processing, including peak detection, noise filtering, feature alignment and data normalization were performed by using the XCMS package in R-project platform. The parameters were applied as follows: the bandwidth was set at 15s, and the peak width was ranged from 5 to 30s; other parameters are selected as default values.

### Statistical Analysis

Univariate analysis was conducted by comparing medians between both participants groups by Mann Whitney test for each metabolite. Multivariate analysis (PCA and PLS-DA plot) was conducted by SIMCA-P v.11.5 (Umetrics, Umea, Sweden) to identify the most influential variables in the differentiation between p7 and p13 with VIP >1.0, fold change >1.5. And cross-validation was repeated 200 times to verify whether the model was over fitted. The heatmap was performed with MetaboAnalyst (http://www.metaboanalyst.ca/MetaboAnalyst/). Cytoscape v.3.1.0 (www.cytoscape.org) was used for correlation network construction. All the statistical analyses for metabolites were conducted by SPSS 19.0 and GraphPad Prism 8.0 software. The data were expressed as mean ± SEM of the three independent experiments.

### Bioinformatics analysis

The sequencing results were compared and identified with reference to the eukaryotic reference system. Differences in gene expression were assessed as fragments per million mapped reads (fpkm) per kilobase of transcript. The screening principle of differentially expressed genes (DEGs) were false discovery rate (FDR) < 0.05, differential expression fold change > 1.5 times. To investigate the biological functions of DEGs, we conducted Kyoto Encyclopedia of Genes and Genomes (KEGG) and Genetic Ontology (GO) analyses for further biofunction enrichment using the WebGestalt database (http://www.WebGestalt.org/).

## Results

### BM-MSCs surface marker expression characterization

Cells were obtained in 2 passages (P2), being continuous cultured, intermediate characterized within 13 passages. Our cells exhibit trilineage mesoderm differentiation ability in 7^th^ passage (Lu et al. 2019). Cells quality attributes were further routinely performed in P7, P9, P11, and P13 (Figure 1A). As shown in figure 1B and figure 1C, BMSCs from developed passage (P13) were much higher in terms of the presence of certain cellular markers, such as CD14 (Magri et al. 2024), HLA-DRB1 (Morita-Fujita et al. 2023) also CD90 (Blair et al. 1997) than from low strains, which indicated an increased tendency in bacterial lipopolysaccharide, viruses, CD4-positive T cells, and be shy of cell communication.

**Figure 1.**
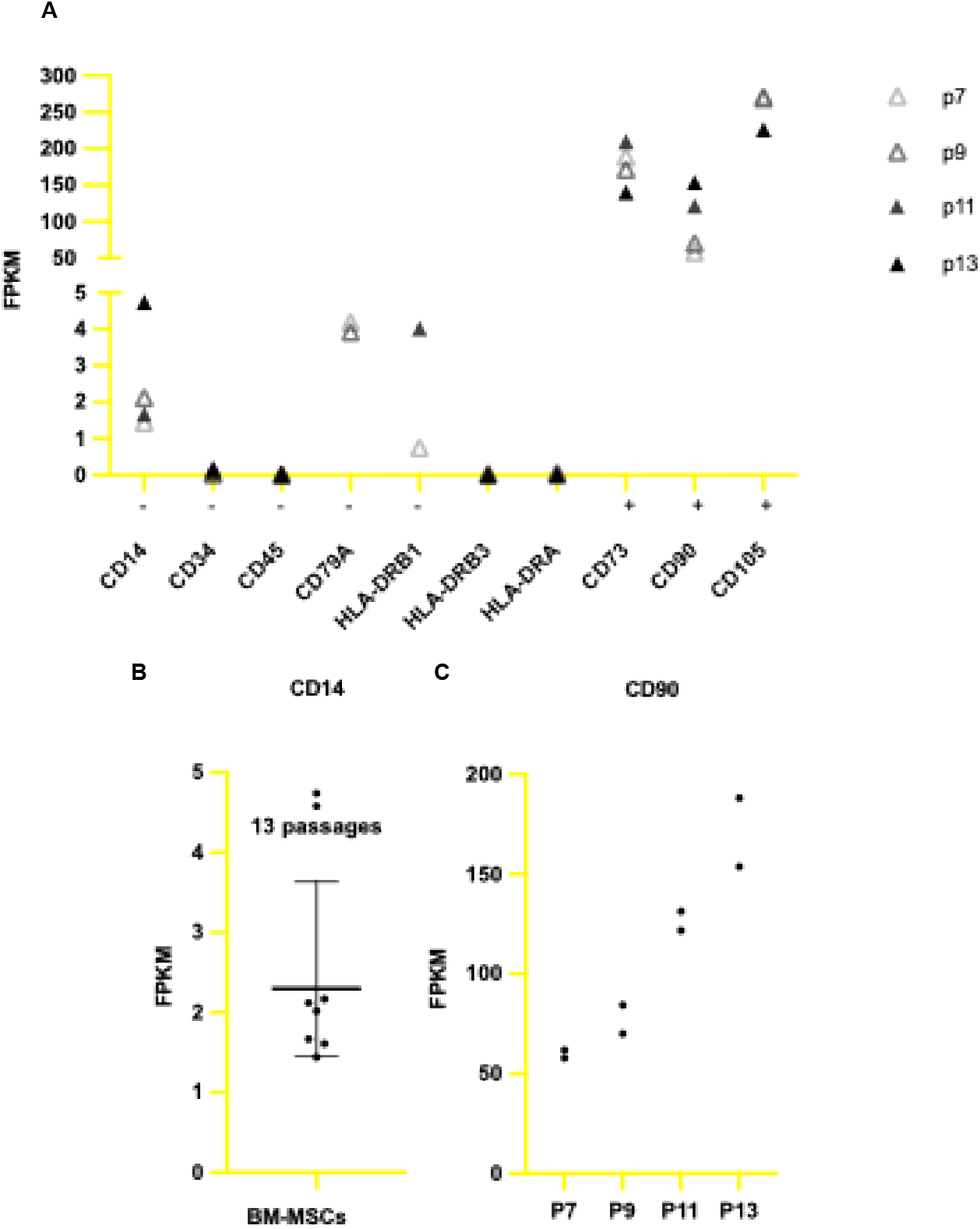
Cell surface marker expression status during passage (P7, P9, P11, P13). A. FPKM of CD14, CD34, CD45, CD79A, HLA-DRB1, HLA-DRB3, HLA-DRA, CD73, CD90, CD105 in P7, P9, P11, P13 BM-MSCs. B. CD14 expression level in invitro cultural BM-MSCs. Geometric mean with geometric SD. C. CD90 FPKM in P7, P9, P11, P13 BM-MSCs.

### PKM and ALDOA enable lactate stasis in cell lysate

BM-MSCs go through continued glycolytic under in vitro keep aging process. As the generations progress and the time passes accomplish cell surface marker disturbance. Enzymes within the cell form the expression pattern of adaptation environmental metabolic influences. Here we report the expression level of whole glycolysis and tricarboxylic acid cycling enzymes. As shown in figure 2, BMSCs establish distinct dynamic enzymatic gene expression signatures. Genes associated with glycolysis (e.g., GPD1, PKM, ALDOA) exhibited increased expression, particularly at later time points, indicating a potential upregulation of glycolytic activity.

**Figure 2.**
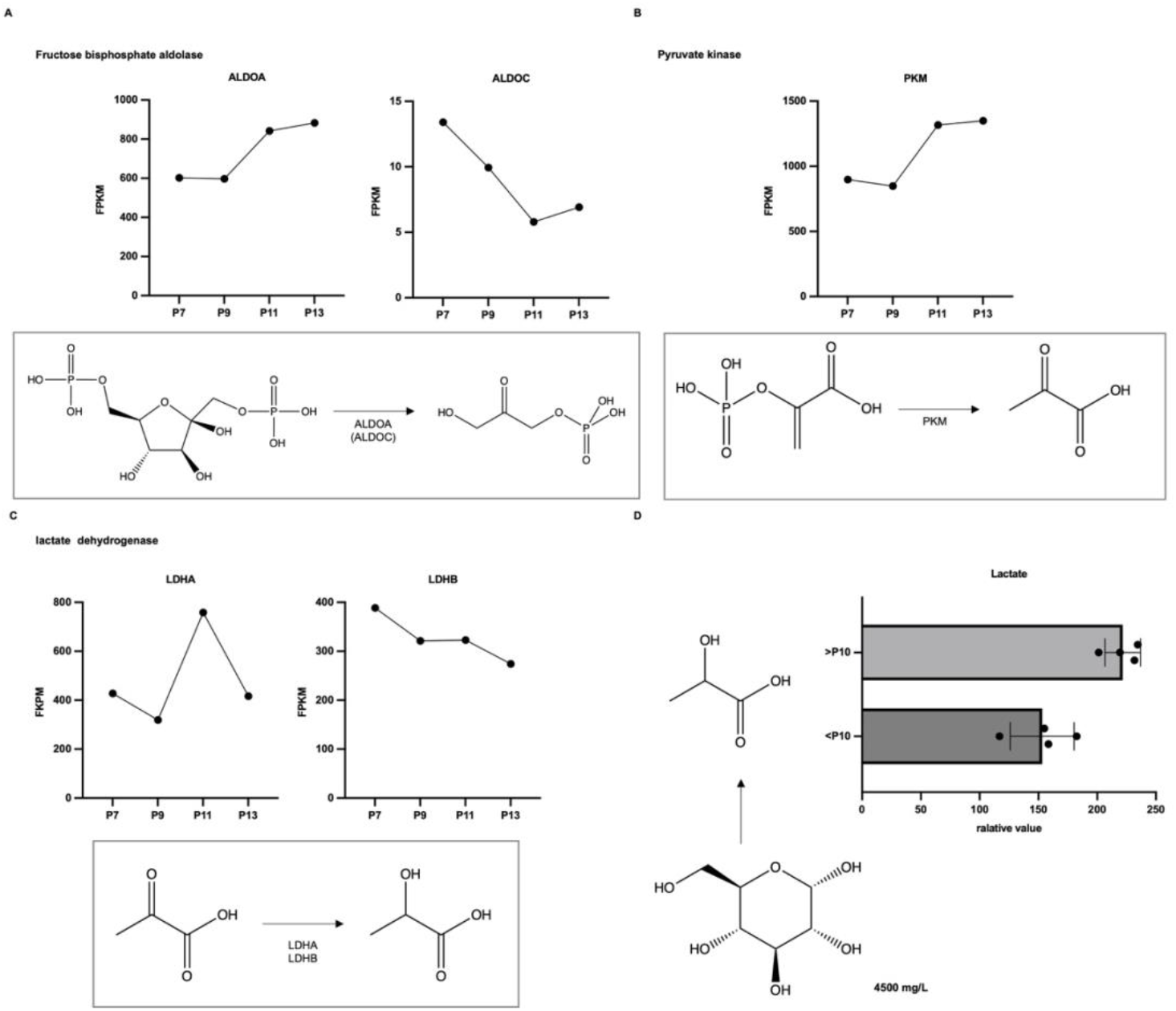
Lactate accumulates in cell cytoplasm. A. Fructose bisphosphate aldolase ALDOA and ALDOC in P7, P9, P11, P13 BM-MSCs. B. Pyruvate kinase PKM expression level in P7, P9, P11, P13 BM-MSCs. C. Lactate dehydrogenase LDHA and LDHB expression level in P7, P9, P11, P13 BM-MSCs. D. Relative value of transformed lactate in P7, P9 BM-MSCs and P11, P13 BM-MSCs. Mean with SD.

### Aging passages BMSCs tricarboxylic acid cycle begin with citrate rise

Cells go through tricarboxylic acid (TCA) cycle, from pyruvate, acetyl-CoA, to citrate, end with malate and oxaloacetate (Figure. 3A), couple with citrate efflux (Figure. 3C) to support cell fit aging process and the requirement medium nutrient components induced citrate accumulation(Xie et al. 2026). Conversely, genes involved in the TCA cycle (e.g., IDH1, IDH2, FH, CS, SUCLG1 and SUCLG2) displayed a consistent decline, suggesting downregulation of mitochondrial energy metabolism. Similarly, genes related to oxidative phosphorylation (e.g., SDH family) showed decreased expression at later stages (Figure. 3B). These findings intended a shift in metabolic activity during cellular senescence, characterized by enhanced glycolysis and suppressed mitochondrial metabolism, with comparable trends observed between the two cell lines. Differences in the magnitude of expression changes reflect age-related differences in metabolic adaptations (Figure. 3C).

**Figure 3.**
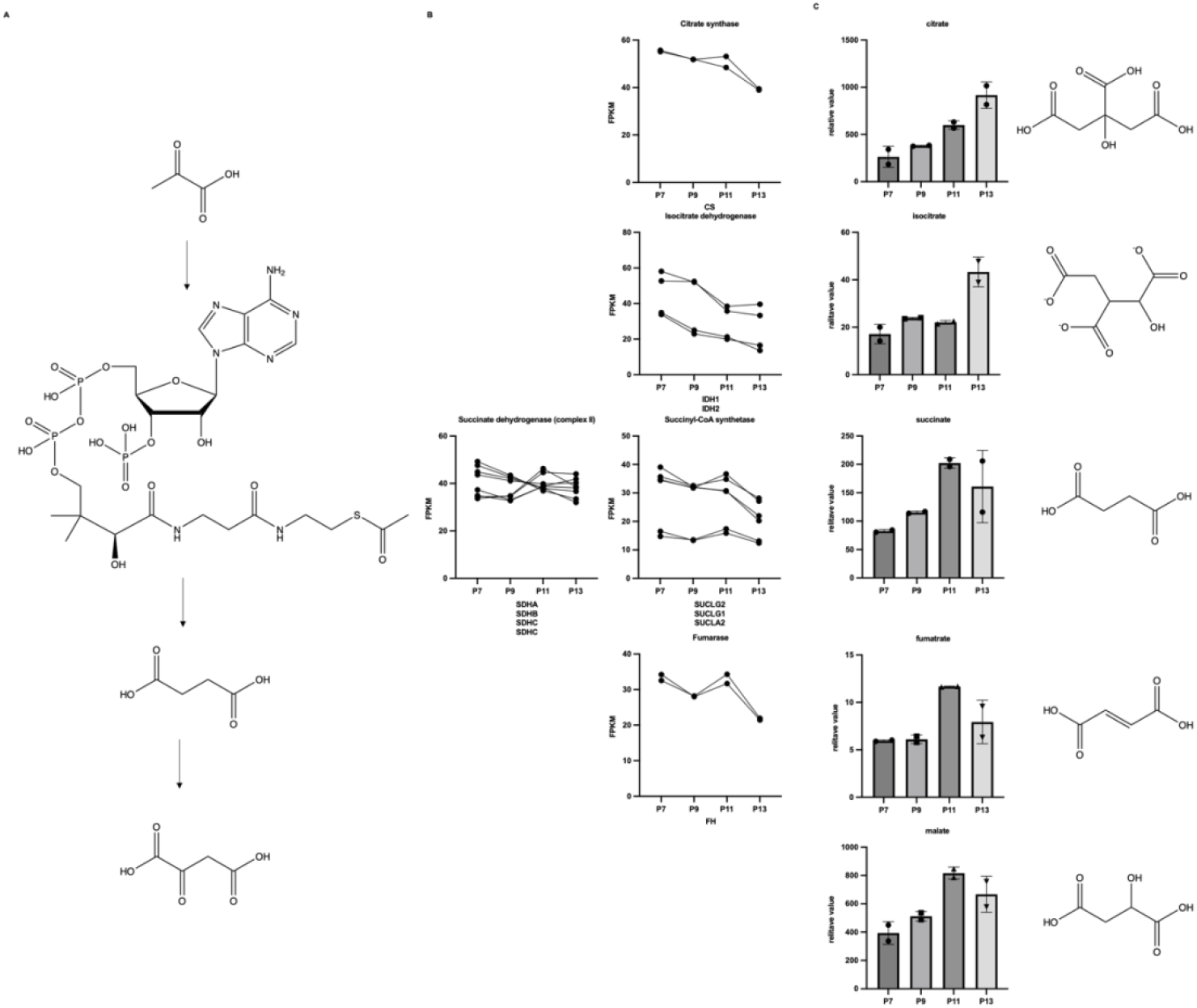
Enzymatic points change within tricarboxylic acid cycle. A. BMSCs main metabolic pathway in TCA. B. The FPKM level of CS (Citrate synthase), Isocitrate dehydrogenase (IDH1 and IDH2), Succinate dehydrogenase (complex II) (SDHA, SDHB, SDHC, SDHD), Succinyl-CoA synthetase (SUCLG2, SUCLG1, SUCLA2) and Fumarase (FH). C. Citrate, isocitrate, succinate, fumarate, malate relative value in P7, P9, P11, P13 BMSCs.

### Mitochondria function

Extracellular acidification rate (ECAR) remained low and stable in p7 cells across all time points (Figure 4a). However, ECAR increased significantly in p13 cells, peaking at mid-time points and declining slightly thereafter (Figure 2a). At the final measurement point, total ECAR was significantly higher in p13 cells compared to p7 cells, indicating a shift toward glycolysis in late-passage cells (Figure 4b). Glycolytic capacity and basal glycolysis were both significantly elevated in p13 cells, further supporting the metabolic transition toward glycolysis with aging (Figure 4c, 2d).

**Figure 4.**
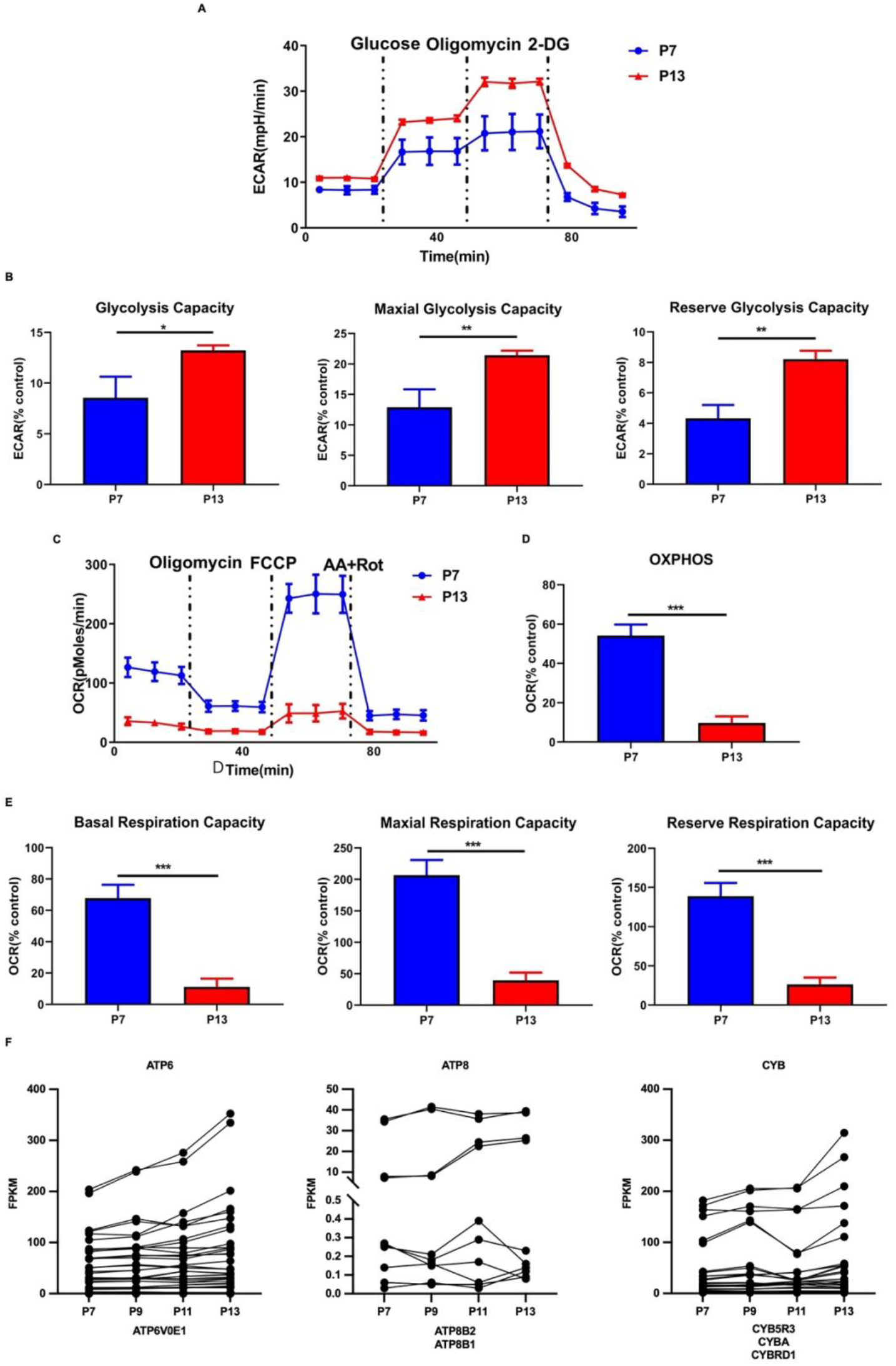
Mitochondrial extracellular flux. A. Glucose metabolism in p7 and p13 cells. The ECAR profiles of p7 and p13 cells following glucose, oligomycin and 2-DG treatments. B. The glycolysis, maxial glycolysis, and reserve glycolysis capacity of p7 and p13 cells. C. Mitochondrial respiration in p7 and p13 cells. D. The OXPHOS capacity of p7 and p13 cells. E. The basal respiration, maxial respiration, and reserve respiration capacity of p7 and p13 cells. F. ATP6, ATP8, CYB FPKM expression level in P7, P9, P11, and P13. *p < 0.05, **P < 0.01, ***P < 0.001.

The oxygen consumption rate (OCR) was measured in early-passage (p7) and late-passage (p13) cells under different treatments. In p7 cells, basal OCR and ATP-linked respiration were significantly higher, peaking at mid-time points and declining at later stages (Figure 3a). FCCP-induced maximal respiration also peaked at mid-time points, indicating robust mitochondrial function in early-passage cells (Figure 3a). In contrast, p13 cells exhibited consistently low basal and maximal OCR across all time points (Figure 3a). At the final measurement point, total OCR was significantly reduced in p13 cells compared to p7 cells, confirming the decline in mitochondrial oxidative phosphorylation with aging (Figure 3b).

Mitochondrial complex activities were also impaired in late-passage cells. Complex I activity was significantly reduced in p13 cells compared to p7 cells, indicating compromised electron transfer (Figure 3c). Complex II and Complex III activities showed similar declines, reflecting broader dysfunction in the electron transport chain during aging (Figure 3d, 3e).

As shown in figure 4f, ATP6: mitochondrially encoded ATP synthase 6, upregulated. Contributes to proton-transporting ATP synthase activity, rotational mechanism. Involved in proton motive force-driven mitochondrial ATP synthesis. Located in mitochondrion. Part of proton-transporting ATP synthase complex.

ATP8: mitochondrially encoded ATP synthase 8, upregulated. Contributes to proton-transporting ATP synthase activity, rotational mechanism. Involved in proton motive force-driven mitochondrial ATP synthesis. Located in mitochondrion. Part of proton-transporting ATP synthase complex.

Mitochondrially encoded cytochrome b (CYB5R3, CYBA) upregulated. This gene is predicted to be involved in mitochondrial electron transport, ubiquinol to cytochrome c. Located in mitochondrion. Part of mitochondrial inner membrane.

ATP synthase positive, also the mitochondrial electron transport process excited, may result in over-polarization.

These findings suggest a shift in metabolic activity during cellular senescence, characterized by enhanced glycolysis and suppressed mitochondrial metabolism, with comparable trends observed between the two cell lines. Differences in the magnitude of expression changes may reflect age-related differences in metabolic adaptations.

### ATP and NADPH

To better explain the adaption of BMSCs accompany the dynamics of medium and cellular aging, whole cell ATP relative value was measured. As shown in figure 5A, with the passage of cells, the relative amount of ATP is increased from P7 to passage 13. To further explore the key metabolism energy substances in BMSCs and exosomes, as shown in figure 5B, ATP, ADP, AMP, GTP, GDP, NADP+, NADH, NAD+ were all in P7 BMSCs, with NADP+/NAD+ 0.4, only NADPH and NAD are detectable in the exosomes of mesenchymal stem cells from umbilical cord, the NADPH/NAD number is 4.2.

**Figure 5.**
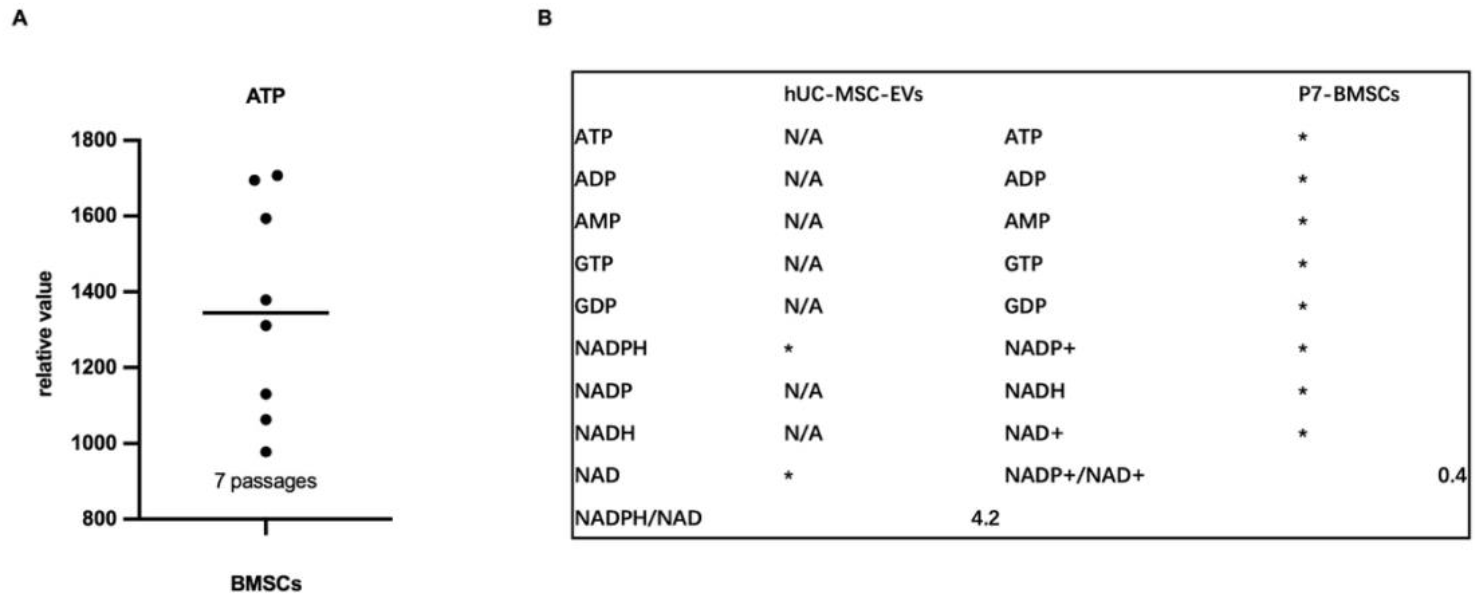
ATPs and NADs family status. A. The relative value of ATP in invitro cultural BM-MSCs. Relative value with median. B. ATP, ADP, AMP, GTP, GDP, NADPH, NADP, NADH, NAD and NADPH/NAD in hUC-MSC-EVs. ATP, ADP, AMP, GTP, GDP, NADP+, NADH, NAD+ and NADP+/NAD+ in P7-BMSCs.

## Discussion

Here, we would like to further discuss the key notes found in our study:

### In vitro passage senescence system

We use consecutive in vitro passage as hBMSCs senescence induction tactics. hESCs can keep undifferentiated status under culture medium over dozens of passage. hBMSCs as adult stem cells, here we change the medium every three days to fresh medium, and passage the cells after they become 70-80% confluence in ratio 1:3. The system is convinced of senescence, ignore the influence of cell secretions, also keep the detection of energy metabolism function in its best condition. At P7, the adipogenic differentiation capacity of cells is significantly reduced, also it is difficult for cells to expand again at P13.

### Mitochondrial respiratory complexes gene changes

Mitochondrial oxidative phosphorylation machinery functions together with the Krebs’ cycle in the matrix. In our research, hBMSCs in vitro passage senescence, three mitochondrial gene expression level changes. ATP8 and ATP6, CYB5R3 and CYBA. A study finds that, co-expressing both ATP8 and ATP6 genes in the nucleus can successful integration into functional Complex V of the oxidative phosphorylation machinery (Boominathan et al. 2016), also achieved substantial success in rescuing a null OxPhos phenotype. Mitochondrially encoded cytochrome b (Feldman and Wainio 1959), a membrane component of the NADPH oxidase complex, handle electrochemical gradient between inner membrane and the intermembrane space/cell cytoplasm via electrons and protons transport.

### Lactic acid excess productive

We measure the total amount of lactate in the cells, we do not characterize the distribution of lactate in each cellular component precisely. Literary research show that lactic acid produces in cell cytoplasm, also can be transitioned to the nucleus and outside the cell through receptors, we did not find any research on lactate entering the mitochondrial membrane, but the increase in intracellular lactate concentration will affect the function of various organelles, it should be a risk for the arrest of P13 cell growth.

### Accumulation of substrates for the tricarboxylic acid cycle

Herein we found each point of tricarboxylic acid cycle general substrates accumulation along enzymic expression level negative regulation accompanies aging. Tricarboxylic acid (TCA) cycle, is broadly conserved from microbes to human beings. The chemical reaction of each turn is condensation of oxaloacetate (OAA), also each turn of the cycle restart with OAA and ends with generation OAA(Arnold and Finley 2023). Four-carbon OAA with two-carbon acetyl-CoA generate citrate, our results show that citrate increased and its synthase decreases, next, isocitrate increases, succinate, fumarate, malate followed by upward trending waves. Isocitrate dehydrogenase (IDH1, IDH2), succinyl-CoA synthetase (SUCLA2, SUCLG2, SUCLG1), fumarase to decay, succinate dehydrogenase (SDHA, SDHD) fades down. TCA itself removes electrons from acetyl-CoA, hBMSCs senescence slowly, TCA sustained generate mitochondrial membrane potential via oxidative phosphorylation (OXPHOS) complex V.

Also, hBMSCs senescence factors exhibit complex interactions between immune regulation and energy metabolism. CD14 influences metabolism through inflammatory responses, HLA-DRB1 affects immune metabolism through antigen presentation, CD90 regulates metabolism in stem cells and the tumor microenvironment, and CD4 impacts overall metabolic states through T cell metabolic reprogramming. These mechanisms may play significant roles in various disease states, providing new targets for disease diagnosis and treatment.

## Data availability

Data will be made available on request.

## Author contributions

All authors shared in the design of the study. All authors wrote the draft of the manuscript, revised it, and agreed with the final version of the manuscript.

